# panomiX: Investigating Mechanisms Of Trait Emergence Through Multi-Omics Data Integration

**DOI:** 10.1101/2025.04.11.648356

**Authors:** Ankur Sahu, Dennis Psaroudakis, Hardy Rolletschek, Kerstin Neumann, Ljudmilla Borisjuk, Axel Himmelbach, Kalyan Pinninti, Nadine Töpfer, Jedrzej Szymanski

## Abstract

Complex omics approaches and high-throughput phenotyping generate large, heterogeneous datasets that make linking molecular signatures to plant traits challenging. To address this challenge, here we introduce panomiX, a user-friendly toolbox for multi-omics integration, designed to enable non-experts to apply advanced computational methods with ease. panomiX automates data preprocessing, variance analysis, multi-omics prediction, and interaction modeling through machine learning, revealing meaningful molecular interactions and synergies. We applied panomiX to a tomato heat-stress experiment combining image-based phenotyping, transcriptomics, and Fourier-transform infrared spectroscopy data, with the aim of identification of condition-specific, cross-domain relationships between gene expression, metabolite levels, and phenotypic traits. Our approach identified a network of such connections, with those linking photosynthesis traits with stress-responsive kinases in elevated temperatures among most significant ones. By simplifying complex analyses and improving interpretability, panomiX offers a platform to accelerate the discovery of trait emergence in plants and select specific candidate genes based on multi-omics analyses.

## Introduction

Recent advances in high-throughput plant phenomics have greatly expanded both the scale and depth of acquired data [1], [2]. Phenomics data are often combined with molecular profiling, including transcriptomics and metabolomics [3], [4], [5], for instance in GWAS experiments [6], time series [7], or genotype-contrast studies [8]. These combined approaches provide valuable insights into the genetic and molecular basis of trait emergence [9], [10]. However, effectively integrating and interpreting such diverse data sets remains a major challenge.

Multi-omics deals with high-dimensional data [11] (**Fig. 1A**) that vary in coverage, variance scales, and exhibit batch effects [12] (**Fig.1B**). Although many specialized tools exist for analyzing specific omics data types (spanning network-based [13], [14], [15], correlation-based [16], similarity-based [17], [18], [19], Bayesian [20], [21], [22], fusion-based [23], and multivariate methods [24], [25], [26]; **Fig. 1C**), these often fail to integrate different omics layers due to usability constraints, complex workflows, or limited interfaces. Consequently, there is an urgent need for a flexible, comprehensive, and user-friendly solution that can seamlessly integrate diverse omics data types using advanced machine learning techniques.

**Fig. 1.**
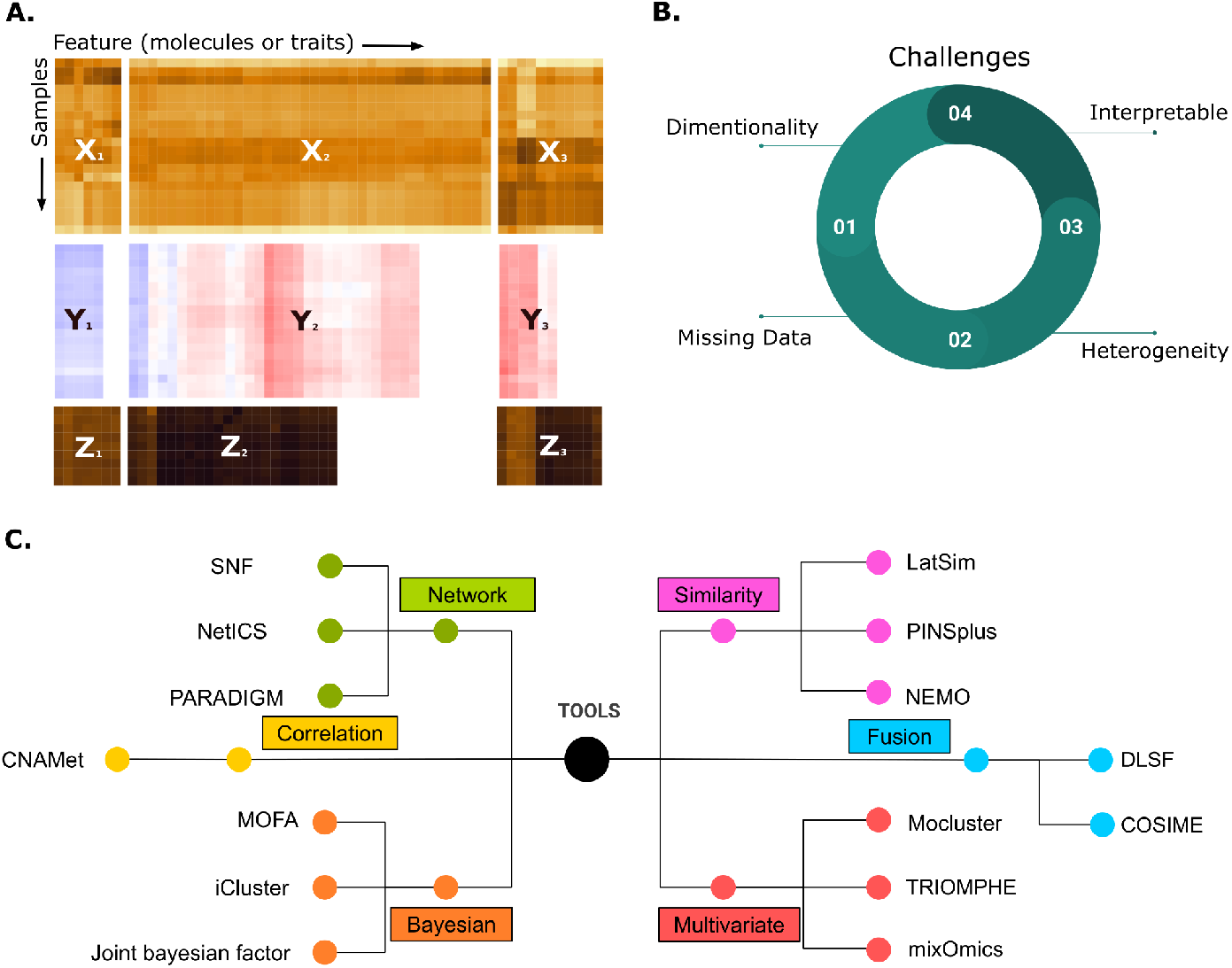
Multi-omic integration - challenges and methods. (A) Omic data sets usually differ in sample size, number of features and variance even if collected in the frame of the same experiment. (B) Different challenges in multi-omics data integration (C) List of available multi-omics data integrative tools.

To address this need, we introduce panomiX, a user-friendly, open-source platform designed for implementing data integration using eXtreme Gradient Boosting (XGBoost) for multi-omic data of varying scales and quality [27]. panomiX employs a harmonization and scaling pipeline prior to data integration, and accommodates both quantitative and categorical data utilizing cross-predictions between omics layers. XGBoost’s inherent ability to handle missing values eliminates the requirement for explicit data imputation. panomiX provides model interpretability using SHapley Additive exPlanations (SHAP) [28], facilitating transparent feature selection, ranking, and visualization. A key feature of panomiX is its capacity to apply user-defined constraints. This enables researchers to test hypotheses on specific biological interactions of interest. This capability extends its utility beyond data integration, allowing for detailed exploration of potential molecular mechanisms between pre-selected sets of phenotype, genotype, and molecular features.

To demonstrate the power of panomiX, we applied it to study the response of tomato seedlings (*Solanum lycopersicum*) to elevated temperatures - an agriculturally relevant process affecting tomato yield both in the greenhouse and field cultivation. Tomato plants are highly sensitive to heat during early growth and later during pollen development, leading to poor germination, inhibited growth, and reduced fruit set [29], [30], [31]. At the molecular level, heat stress induces transient gene reprogramming, triggering heat shock proteins (HSPs) and heat shock factors (HSFs), often regulated by MAP kinases and Cysteine-rich Receptor-like kinases (CRKs) [32], [33], [34], [35], [36]. Multi-omic approaches capturing transcriptomic, metabolomic, and protein data were crucial to unraveling some elements of these complex regulatory networks [37].

In this study, we analyzed the heat stress response and subsequent recovery of young tomato seedlings under controlled conditions, combining daily high-throughput phenomic profiling with RNA-seq analysis and Fourier-transform infrared spectroscopy (FTIR) at selected time points. We describe both common response patterns across all data types and those unique to specific datasets. Further, we highlight key interactions among transcripts, metabolic features, and phenotypes uncovered by panomiX and link these to relevant biological processes. Throughout, we demonstrate the functionality and versatility of panomiX for multi-omics data integration and analysis.

## Materials and Methods

### Data preprocessing and analysis

The panomiX platform integrates several specialized R libraries for data harmonization, variance analysis, multi-omics prediction, interaction, and visualization. We provide default data preprocessing methods for transcriptomics and FTIR data. For transcriptomics, we provide a ‘DESeq2-based’ [38] approach, adjusting for factors like sequencing depth and sample composition. For FTIR spectral data, the ‘baseline’ R library [39] is employed to remove background noise, and the ‘signal’ library [40] for Savitzky-Golay smoothing [41] of spectral features without loss of peak sharpness and spectral integrity. Variance analysis is conducted using the ‘irlba’ R library and prcomp function [42], including dimensionality reduction, clustering, and identification of sources of variance. Multi-omics prediction workflows are facilitated by ‘caret’ R library [43], with the built-in framework for hyperparameter tuning and model evaluation. XGBoost is used for predictive modeling, and SHAP and Boruta-SHAP are employed for model interpretability and feature selection [44].

### Visualizing insights with interactive plots

The panomiX user interface (UI) uses ‘shinyjs’ [45] to create dynamic interactions, allowing users to control UI element visibility based on their actions, for a streamlined experience. Interactive plots are created with ‘plotly’ [46], enabling users to zoom, pan, and hover over data points for deeper exploration. The ‘DT’ package [47] enables interactive data tables, facilitating sorting and exploration of large datasets. ‘shinycssloaders’ [48] provides visual feedback on application progress during data loading. Additionally, ‘bslib’ [49] offers advanced plot formatting tools, and ‘bsicons’ [50] incorporates intuitive icons, enhancing UI usability and aesthetics.

### Source Code Availability and Community Collaboration

The panomiX toolbox is developed fully in R [51] and Shiny [52] and deployed on Shinyapps.io https://www.shinyapps.io/. The source code is managed with a GitHub repository connected to the Shinyapps.io via ‘rsconnect’ [53]: https://szymanskilab.shinyapps.io/panomiX/. The source code for the platform is available on GitHub: https://github.com/NAMlab/panomiX-tool. The repository contains all the necessary R scripts for data processing, visualization, and machine learning prediction.

### Experimental data and design

Seeds of genotype *Moneymaker* were planted in well-watered soil trays and kept under a plastic cover in a walk-in phytochamber with a day/night cycle of 16/8h and temperatures of 24°C and 20°C respectively. The seeds were kept without light for one day, then illumination of 320µmol per square meter per second (from from Whitelux Plus metal halide lamps, Venture Lighting Europe Ltd., Rickmansworth, Hertfordshire, England) was added during the day phase. After a total of 7 days after sowing, evenly germinated and healthy seedlings were transplanted into individual pots (10 cm diameter, 8 cm height) without plastic covers at 60-70% relative air humidity. After 9 further days of establishment in the pot, the plants were exposed to heat stress at 37°C/28°C (day/night) for 6 days followed by a recovery phase at 24°C/20°C. Plants were watered once or twice daily by an automated system replenishing evapotranspirated water by weight to ensure the plants do not experience drought stress. All plants were phenotyped daily using an imaging-based high-throughput system and the youngest fully formed leaf pair was harvested (directly put in liquid nitrogen and then stored at -80°C) for molecular measurements mid-day (between 13:00 and 14:00, light phase was from 06:00-22:00) from individual plants directly before the temperature switch, each day during the heat stress phase and after 1, 2, and 4 days of recovery. Control plants were grown exactly the same way except that they stayed at 24°C/20°C throughout the whole experiment. They were sampled at the same respective time points as the heat-treated plants.

### Data acquirement

During daily phenotyping, plants were photographed from the top and three side angles in visible and in fluorescent light (excitation: 400-500 nm, emission: 520–750 nm). The images were then analyzed using the Integrated Analysis Platform (IAP) software [54] to yield 138 phenotypic traits for each measurement, as described in [55]. Next we reduced the 138 phenotypes into a smaller set of non-correlated variables. Using hierarchical clustering, we grouped related phenotypes together, and each group was represented by its first principal components (PCs). For each group, we evaluated the explained variance of the first PC and applied a threshold of 50% explained variance as the cutoff. Through this process, we narrowed the phenotypes down to 27 key representatives, which we used for model training.

For the molecular assays, the frozen leaves were ground and aliquoted for specific measurements: Total RNA was isolated from 70mg of the ground material using the RNAeasy Plant Mini Kit (QIAGEN) according to the manufacturer’s protocol. The construction of sequencing libraries involved the Illumina stranded mRNA Prep Ligation Kit (standard Illumina protocol; Illumina, San Diego, California, USA) and 1 µg DNAse I digested total RNA. Sequencing of equimolar library pools (average size: 332 bp) was performed on an Illumina NovaSeq 6000 device (IPK-Gatersleben), using the XP-workflow and a S2 flowcell (Illumina, San Diego, California, USA). On average, 41 M reads (single reads, 118 cycles) were generated per sample. Reads were then mapped to the ITAG4.1 tomato reference genome (https://solgenomics.net/ftp/tomato_genome/annotation/ITAG4.1_release/) using our rnaseq-mapper pipeline built around kallisto [56], available at https://github.com/NAMlab/rnaseq-mapper; [57].

FTIR analysis of freeze-dried leaf material was performed as in [58]. Briefly, approximately 2 mg of freeze-dried material was used per sample. Spectra were produced from ATR-FTIR measurements using the INVENIO-S FTIR spectrometer (Bruker Optics, Ettlingen, Germany) with a Globar light source under continuous purging with dry air. ATR absorbance spectra were recorded in the spectral range of 4000-400 cm-1 at a spectral resolution of 4 cm. Each spectrum consisted of 32 co-added scans. As a background reading, the spectrum of the empty ATR crystal was collected prior to measurement and subtracted automatically from each recorded spectrum using the OPUS software (Bruker Optics).

## Results

### Data input

Our experiment provided 138 phenotypic variables reduced to 27 uncorrelated phenotype clusters (**Fig. S1**), 34688 gene expression values, and 2526 FTIR data points across 10 time points and 2 to 3 biological replicates in two experimental conditions: control growth, and treatment with 6 days of heat stress treatment followed by 3 days recovery phase (**Fig. 2**).

**Fig. 2.**
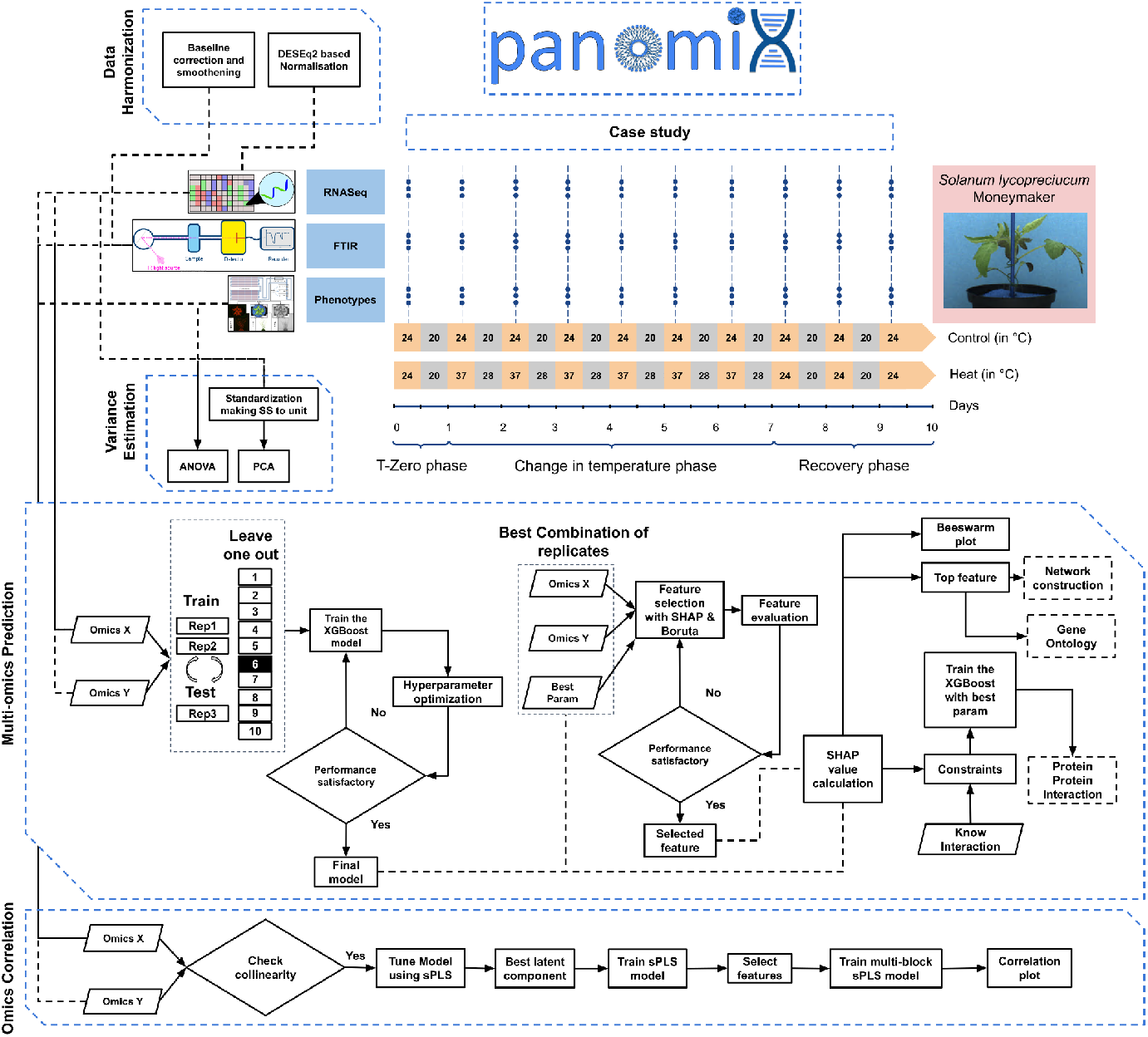
Flow diagram of the panomiX toolbox and the experimental design for the tomato heat stress case study.

#### Tool settings and recommendations

PanomiX works with continuous molecular data, such as normalized RNA-seq counts, protein abundances, metabolite concentrations, or FTIR spectra. Your data should be in a feature matrix format, where: Columns represent biological samples (e.g., individuals or time points). Rows represent molecular features (e.g., genes, proteins, metabolites, or spectral variables). A continuous outcome variable (y) is needed for regression tasks. PanomiX can handle large datasets with tens of thousands of features, but pre-filtering your data is recommended for better performance and computational efficiency. Here’s how you can optimize your dataset: filter low-variability features, remove features that show little variation across samples, exclude low-count features for sequencing data (like RNA-seq), and drop features with consistently low counts, remove near-zero variance features which contribute little to the model and can be safely excluded. These steps help reduce the size of high-dimensional datasets, minimize overfitting, and improve the speed of machine learning algorithms like XGBoost during model training and hyperparameter tuning. Before using panomiX, ensure that your datasets are pre-processed and normalized according to the requirements of the specific ‘omics platforms.

### Data processing and filtering

The RNA-seq data processing with rnaseq-mapper [https://github.com/NAMlab/rnaseq-mapper] provided a data matrix of estimated TPM values (transcript per million, kallisto estimates; [56]) for 49 samples (both control and treatment), with columns representing samples and rows representing genes-aggregated transcript levels. We used TPM counts without applying the data harmonization component for further analysis. To filter out transcripts with consistently low TPM counts, we retained only those transcripts that have at least one sample with a count of 50 or higher, ensuring that transcripts with TPM counts below 50 across all samples were removed. Ultimately, we used 5,479 transcripts for further analysis. Similarly, for FTIR spectral data, we created a data matrix of absorbance values, containing 49 samples (both control and treatment). We selected the absorbance values in the range from 4,000 to 400 cm−^1^. We then applied FTIR normalization using the data harmonization component in the panomiX toolbox. For illustration **Fig. 3A** presents the raw FTIR spectra before preprocessing. After baseline correction and Savitzky-Golay smoothing, the normalized spectra are displayed in **Fig. 3B**, illustrating the improved signal quality.

**Fig. 3.**
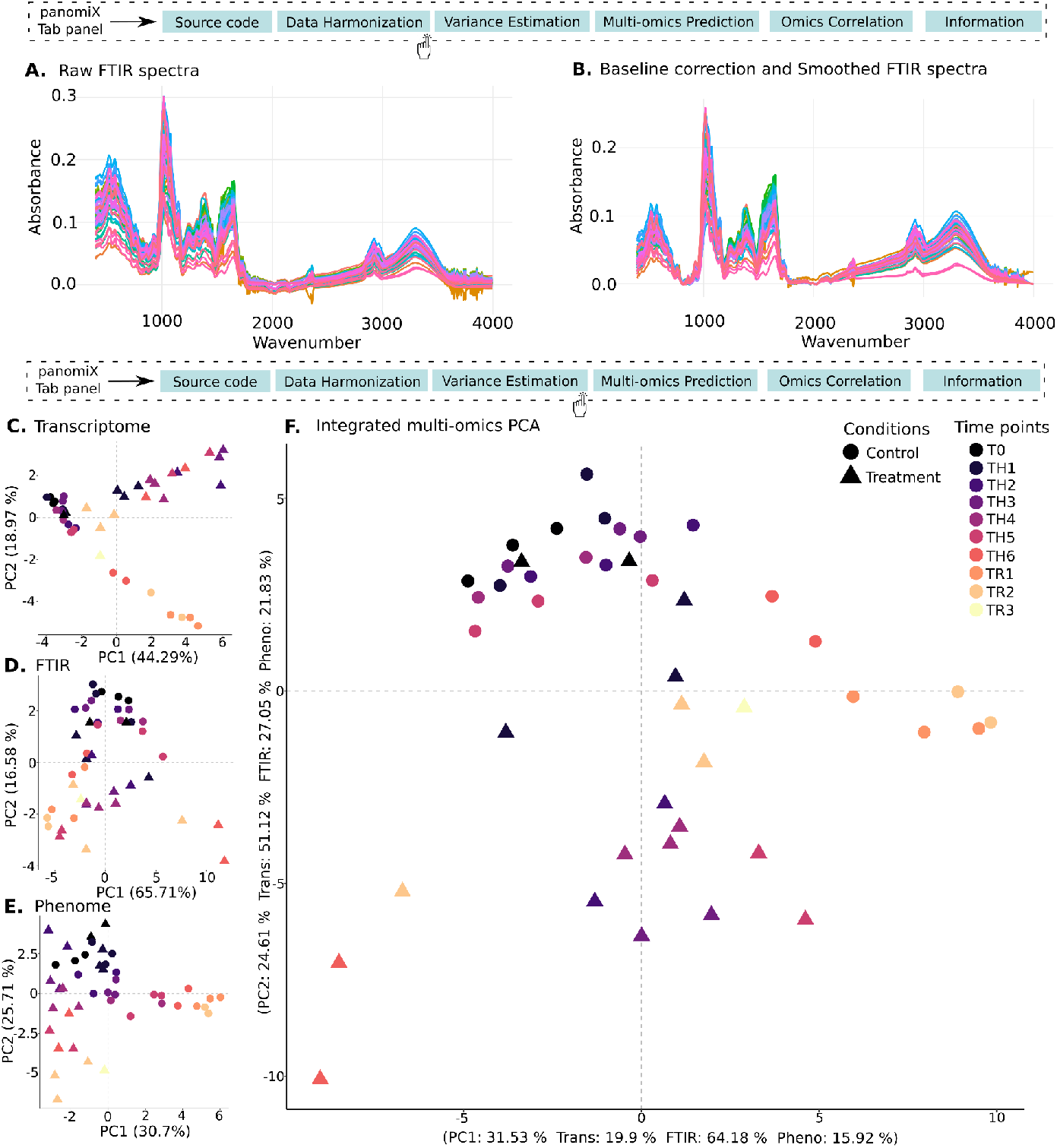
Data harmonization and variance estimation component (A) Raw FTIR spectra (B) Baseline corrected and smoothed FTIR spectra. Individual PCA for each omics data (C) Transcriptome, (D) FTIR, and (E) Phenome. (F) An integrative PCA, where the multi-omic data matrix was combined to identify shared variance components.

#### Tool settings and recommendations

panomiX provides standard methods for normalization of transcriptomic data. First, a log transformation is applied to stabilize variance and improve comparability. Then, the transformed gene expression data is used to determine the geometric mean for each gene. Next, genes with an infinite geometric mean are filtered out, allowing us to focus only on housekeeping genes. The previously log-transformed value of each gene is then subtracted from the respective geometric mean. This step helps identify genes within each sample that have higher or lower expression levels than the average. Subsequently, the geometric mean-subtracted values are used to determine the median, which helps reduce the influence of high-value outliers. Since genes with higher expression values are fewer in number, the median primarily reflects the housekeeping genes. Finally, the median values are transformed back from log to normal values to obtain the final scaling factor. The raw read counts are then divided by these scaling factors. This log and median-based scaling method effectively corrects batch effects and extreme variations. For consistency, we recommend performing all steps of data normalization from the raw counts within panomiX. However, the platform also accepts normalized transcriptomics data in TPM or CPM counts. For the spectral data, such as those obtained by FTIR, panomiX utilizes the baseline correction on the raw spectrum to remove artifacts and the background noise. Following this, Savitzky-Golay smoothing can be applied to each baseline-corrected spectrum. This step applies a polynomial smoothing filter to reduce high-frequency noise while maintaining essential spectral features.

### Variance analysis

The exploratory analysis of the multi-omic data variance highlighted both data type-specific, and shared patterns of changes driven by plant development and heat-stress treatment. In the transcriptome PCA (**Fig. 3C**), PC1 (44% variance) largely reflects time-dependent changes in gene expression, while PC2 (18% variance) highlights heat-stress effects. Early-stage controls and T0 heat samples cluster together, indicating minimal transcriptional differences at the start, whereas later-stage heat treatments diverge, revealing progressive stress adaptation. In FTIR absorbance data (**Fig. 3D**), PC1 (65% variance) again captures temporal shifts, and PC2 (16% variance) distinguishes stress from control conditions; notably, mid-stage heat-treated seedlings cluster with recovery-stage controls, indicating partially overlapping metabolic responses. Phenomic data (**Fig. 3E**) show stronger treatment separation on PC1 (30% variance) and time progression on PC2 (25% variance), where early-stage controls resemble T0 heat samples, and later stages separate more distinctly. Integrating all omics layers (**Fig. 3F**) confirms the interplay of time and treatment: PC1 (31% variance) is dominated by FTIR and transcriptome signals, clearly separating recovery-stage heat-treated seedlings, while PC2 (24% variance), driven by transcriptional variation, further distinguishes controls from heat-stressed plants at various stages. These results underscore the distinct yet complementary biological insights gained by analyzing temporal and stress effects across transcriptomic, metabolic, and phenotypic data.

#### Tool settings and recommendations

PanomiX supports exploratory data analysis with PCA or ANOVA to identify dominant sources of variance. Before PCA, all datasets are standardized to a unit sum of squares, ensuring comparable total deviations across different data types, but all the relevant dataset-specific normalization steps should be performed beforehand. First, PCA is performed separately on each omics dataset to capture dataset-specific variance patterns. Second, the integrated PCA is conducted on a combined dataset to reveal shared variance and global patterns. For PCA, the tool automatically handles data centering and scaling. For the ANOVA and plotting the PCA scores, the user needs to upload metadata with at least an ID column (matching the omics data) and respective condition columns if experimental factors are present. PanomiX often exposes differences in resolution and variability among different omic datasets, demonstrating how multi-omics integration can also provide insights into each individual dataset. Notable, these differences should be carefully evaluated, as differences in normalization, coverage and data filtering procedures can confound their biological importance.

### Multi-omics prediction components

Linking the high-throughput profiling of plant traits with molecular assays such as RNA-seq or metabolomics, can yield valuable biological insights into molecular mechanisms of trait emergence. To explore these connections, we conducted cross-prediction on phenotypes using transcriptomics alone, FTIR alone, and combined transcriptomics + FTIR data, each trained separately on control and treatment samples. This approach enabled identification of condition-specific molecular markers. Model performance, assessed via mean R-squared values across three replicate splits, showed generally higher accuracy in control samples; however, some treatment-based models (e.g., Phenol–Chlorophyll Ratios (side), Mean Fluorescence Intensity (side)) outperformed their control counterparts (**Fig. 4A and 4B**). Next, we performed gene ontology (GO) enrichment on transcripts with high SHAP values (i.e., highly predictive of phenotype), focusing on models with R^2^ > 0.5 and replicate-model R^2^ > 0.7. This criterion yielded 10 control and 7 treatment models (**Fig. S2A and S2B**). Certain phenotypes had unique GO enrichments, while others overlapped. For instance, a control model predicting Relative Fluorescence Area Change (side) highlighted Glutamate decarboxylase and Sugar transporter ERD6-like 6 both linked to enhanced photosynthetic capacity [59] (**Fig. 4C**). Meanwhile, the corresponding treatment model identified cytochrome b559 subunit alpha, part of the photosynthetic electron transport chain (Photosystem II), and phylloplanin, which contributes to stress defense via type VI glandular trichomes [60], [61] (**Fig. 4D**). Next, we extended the analysis to FTIR and combined transcriptomics+FTIR models, applying the same performance thresholds (R^2^ > 0.5 overall and R^2^ > 0.7 for individual replicate models). This revealed a set of FTIR spectra closely linked to each phenotype. Notably, several transcripts and FTIR features were consistently identified as top predictors across all three omics-specific models (**Table S2**). For example, in the Relative Fluorescence Area Change (side) phenotype (**Fig. S3**), the transcript-only model pinpointed cytochrome b559 subunit alpha and phylloplanin as key contributors (**Fig. S3A**). In the FTIR-only model, spectra 2002 and 2003 showed strong predictive power (**Fig. S3B**). The integrated model (transcriptomics+FTIR) retained these same predictors (**Fig. S3C**), confirming their importance. Further analysis of the transcript-to-FTIR prediction revealed that phylloplanin significantly predicted the 2002 FTIR spectrum (**Fig. S3D**).

**Fig. 4.**
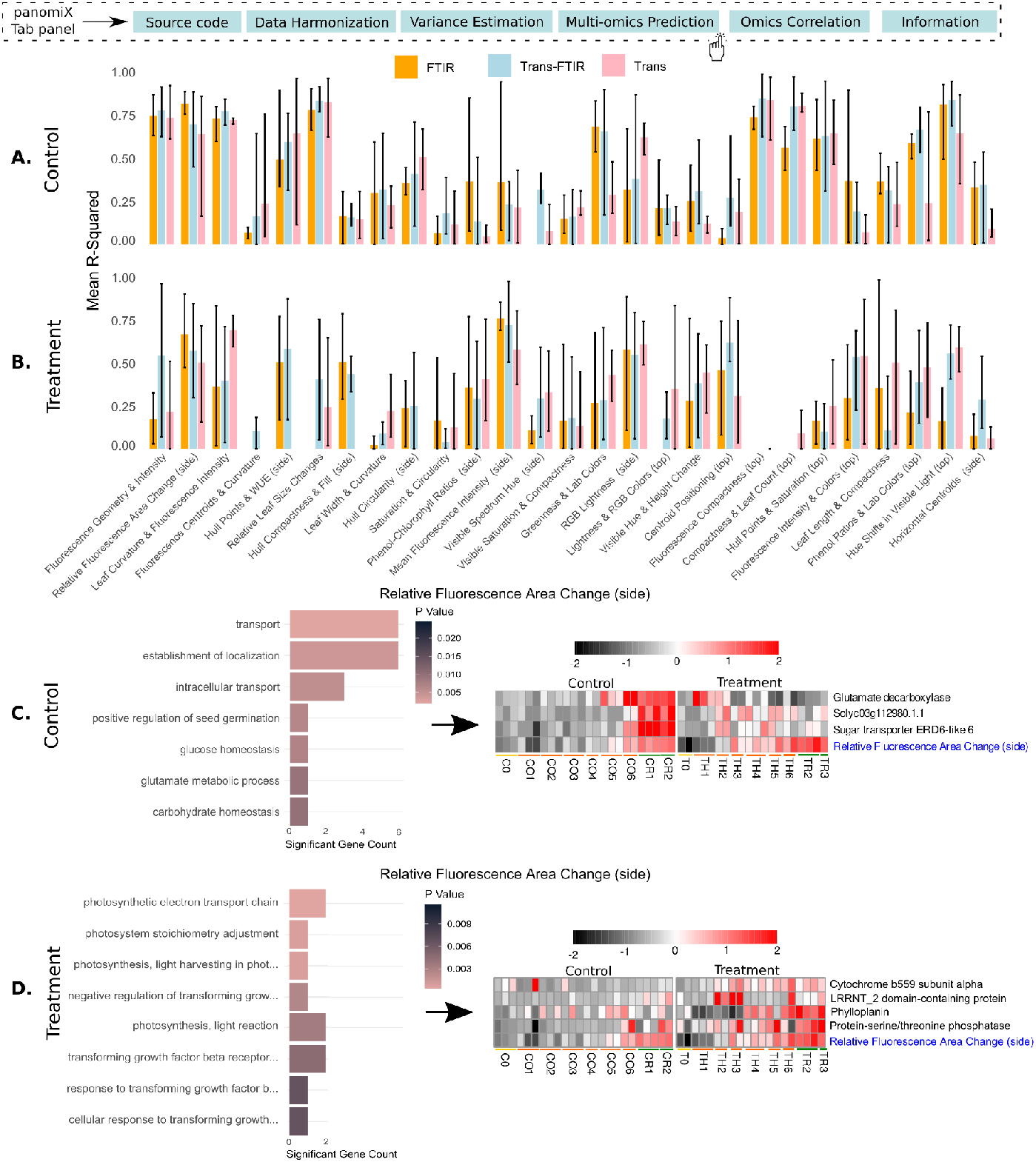
(A) Performance of control samples for transcripts predicting phenotypes, FTIR predicting phenotypes and combined transcripts and FTIR predicting phenotypes (B) Performance of treatment samples for transcripts predicting phenotypes, FTIR predicting phenotypes and combined transcripts and FTIR predicting phenotypes (C) Gene ontology and expression profile for the features selected for Relative Fluorescence Area Change (side) from the control model (D) Gene ontology and expression profile for the features selected for Relative Fluorescence Area Change (side) from the treatment model.

#### Tool settings and recommendations

Training the XGBoost model requires splitting the data into training and test samples. Users can split datasets either randomly (using a training size slider) or by replicate to ensure consistent grouping of the same replicates, and hyperparameter tuning is automated via the caret method. In the cloud version of panomiX, two main hyperparameters (number of rounds, tree depth) can be configured, with others (learning rate, gamma, subsample, etc.) preset to optimal ranges. The desktop version provides greater flexibility, letting users adjust a wider range of parameters (see the full documentation; https://github.com/NAMlab/panomiX-tool). Model generalization is validated through cross-validation (CV) or leave-one-out cross-validation (LOOCV), and performance is reported using R^2^ and RMSE. A model performance plot (R^2^) is displayed, alongside a feature-importance table that ranks predictors. SHAP values further reveal how individual features positively or negatively influence phenotype predictions, visualized via beeswarm plots for intuitive interpretation. The Boruta-SHAP algorithm is also available for alternative feature selection, with its results shown in a separate beeswarm plot. For biological replicates, we recommend replicate-based splitting (with metadata specifying sample names and replicate identifiers) to improve model consistency and accuracy. Random splitting with an adjustable training size can be used for general evaluation. In the cloud version, tuning the number of rounds and tree depth is crucial for better performance; the desktop version allows deeper parameter control. CV is advisable for robust model generalization, while LOOCV suits smaller datasets. For deeper insights, users can leverage SHAP values and feature-importance rankings to interpret and prioritize key predictors.

### Multi-omics prediction with interaction constraints

PanomiX can analyze interactions of predictive features identified by the model as well as “known” features provided by users - such as genes of interest - by incorporating both sets into a single prediction framework. It uses SHAP values to show how these features (positively or negatively) influence a given outcome (e.g., a phenotype) and to set constraints for the model (**Fig. 5A**). In our experiment, these constraints relate to a predetermined list of literature- and annotation-based candidate transcripts. We tested two scenarios: a) evaluation of data-derived candidate transcript set; and b) evaluation of a predetermined list of candidates.

**Fig. 5.**
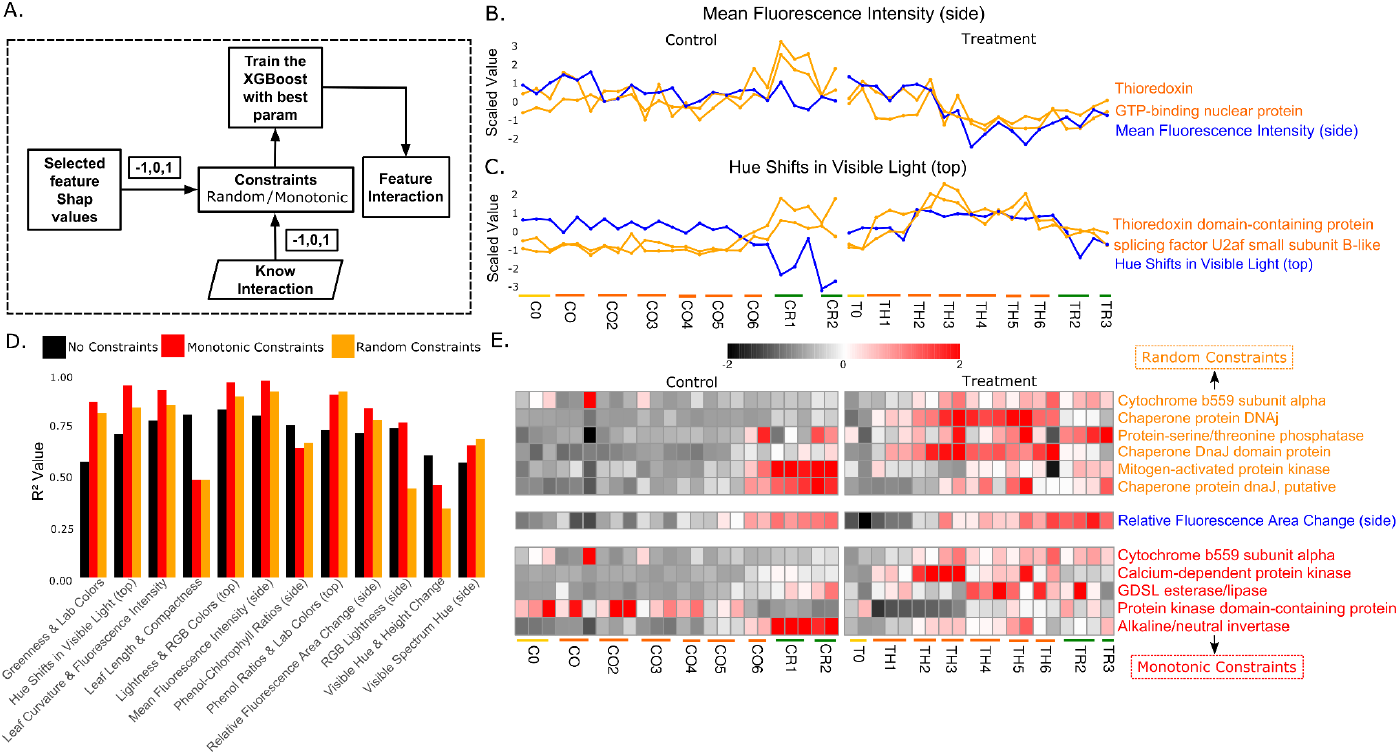
Feature interaction. (A) Illustration of feature interaction as a flow diagram (B) Interaction between the heat stress related (scenario a) transcripts and predictive features from trained model Mean Fluorescence Intensity (side) (C) Interaction between the heat stress related (scenario a) transcripts and predictive features from trained model Hue shifts in Visible Light (top) (D) Performance of treatment samples for transcripts predicting phenotypes (scenario b). Here Black color bar represents the model trained without constraints, the red color bar represents the model trained with monotonic constraints, and the orange color bar represents the model trained with random constraints (E) Expression profile for the features selected from the constraints model of Relative Fluorescence Area Change (side) (scenario b). Here orange colored features are selected from the random constraints model and red colored features are selected from the monotonic constraints model.

In the first scenario, we tested 57 heat-stress-related transcripts (46 up-regulated, 11 down-regulated; Dennis et al. [57]) alongside the features previously identified by panomiX. Notably, several transcripts showed positive interactions with both the predicted features and the target phenotypes (**Table S3**). For instance, in a heat-treatment model predicting Mean Fluorescence Intensity (side), GTP-binding nuclear protein (down-regulated under heat stress) maintained its down-regulation relative to controls, aligning with its predictive role. Similarly, in a model predicting Hue Shifts in Visible Light (top), thioredoxin domain–containing protein interacted with splicing factor U2af small subunit B-like, suggesting a joint effect on the phenotype under heat stress (**Fig. 5B and 5C**).

In the second scenario, we explored whether stress responses extend beyond heat shock proteins to kinases and other signaling elements. For that purpose, we compiled 1,531 candidate genes (99 guanylate cyclases, 1,038 other kinases, 394 heat-related genes) from literature and databases [62], [63], [64]. We then trained models using no constraints, random constraints, or monotonic constraints (**Fig. 5D**). Most cases showed improved performance with random constraints over no constraints, and an additional ∼5% improvement under monotonic constraints. A representative example is a model predicting Relative Fluorescence Area Change (side), where random constraints boosted performance by 5% compared to no constraints. Key transcripts (e.g., cytochrome b559 subunit alpha, protein-serine/threonine phosphatase) interacted with MAP kinase and DnaJ domain proteins (**Table S4**). Under monotonic constraints, performance rose another 5%, and cytochrome b559 subunit alpha was linked to GDSL esterase/lipase and calcium-dependent protein kinase, both implicated in stress response [65], [66], [67].

Overall, these findings highlight panomiX’s capacity to reveal significant interactions among user-specified and model-predicted features, offering mechanistic insights into phenotype determination and guiding further experimental validation.

#### Tool settings and recommendations

By including a user-provided list of relevant (or potentially important) known features, PanomiX can determine whether these inputs significantly contribute to the model’s outcome or interact with features already identified by the model. The “already predicted features” are those uncovered after training, together with their SHAP values, which may be positive or negative in relation to the outcome. PanomiX uses these SHAP values to set constraints, illustrating each feature’s relationship with the predicted outcome. In ‘random constraints’ setting, the direction (positive or negative) of feature-outcome relationships is not predefined. In this case, users provide a table with two columns—”feature” and “final_association”—where “final_association” contains only 0 (meaning the relationship is unknown). PanomiX then applies random constraints using this information together with the “predicted outcome.” In the ‘constrained’ setting, monotonic constraints require a table with the same columns, but users assign 1 for a known positive relationship, -1 for a known negative relationship, or 0 if it is unknown. PanomiX merges this information with the “predicted outcome” from the trained model, ensuring predictions remain consistent with domain knowledge.

### Multi-omics prediction components vs mixOmics

To evaluate the efficiency, explainability, and stability of the panomiX multi-omics cross-prediction functionality, we tested it against mixOmics [25], a comparable state-of-art tool offering similar functionalities. Specifically, we compared the statistical methods offered by both tools as a main technique of data integration, namely the sparse Projection to Latent Structures (sPLS) of mixOmics with the XGBoost implemented in panomiX. For that purpose, we conducted two cross-prediction analyses: a) transcriptomics data was used to predict phenotypic traits; b) the same phenotypic traits were predicted from the FTIR data (**Fig. 6A and 6B**).

**Fig. 6.**
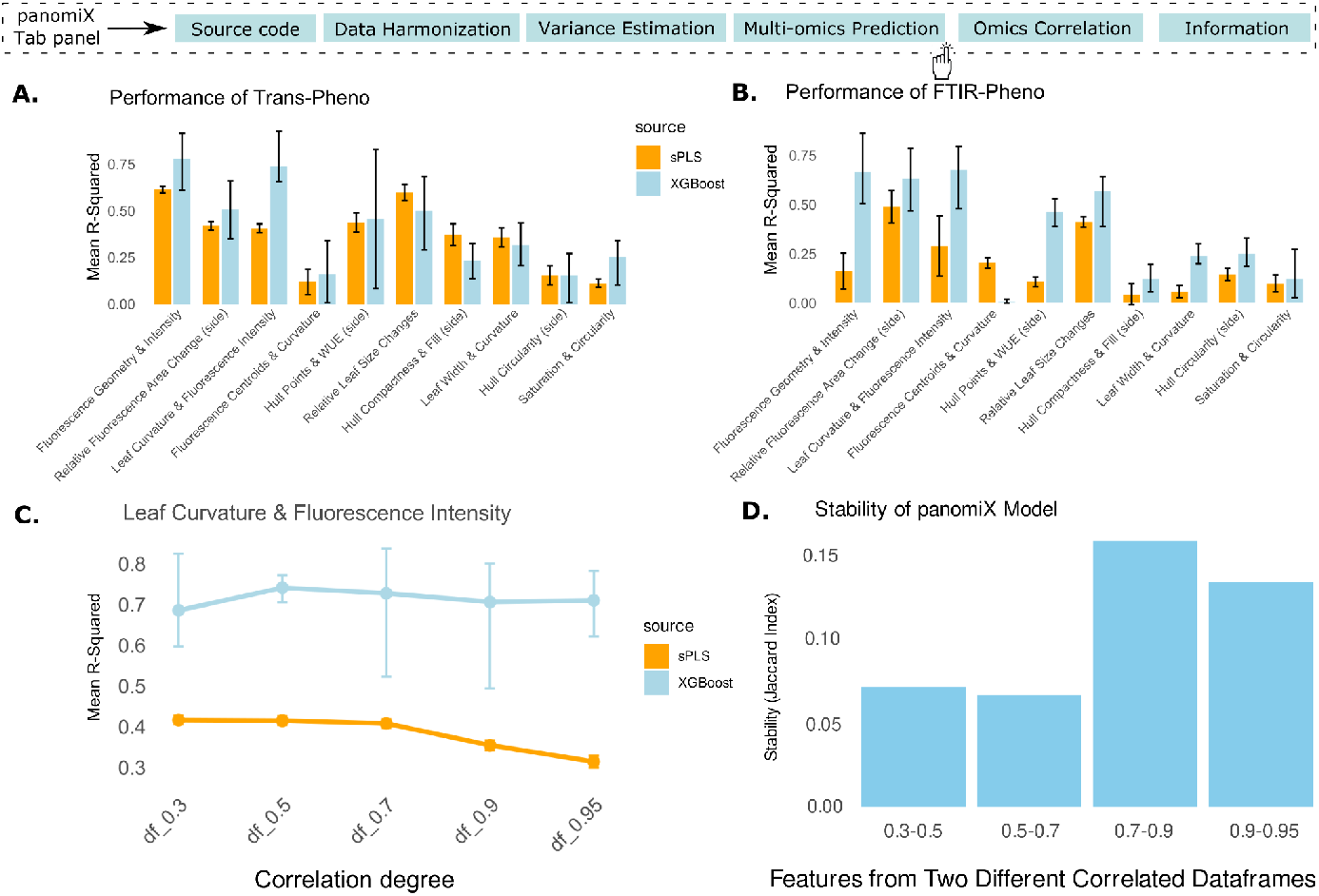
Multi-omics integration in panomiX and mixOmics. (A) Performance for transcripts-phenotype prediction. (B) Performance for FTIR-phenotype prediction. (C) Effect of multicollinearity on model performance. (D) Robustness of feature selection for different thresholds of the feature collinearity in the panomiX XGBoost implementation.

To further investigate robustness, we compared panomiX and mixOmics (sPLS) on transcriptomics subsets with varying correlation coefficients (0.3 to 0.95), using one phenotype cluster (Leaf Curvature & Fluorescence Intensity) as a test. Despite sPLS’s known strength with highly collinear data, panomiX consistently outperformed it (**Fig. 6C**). We also evaluated feature importance stability across the correlation gradients by comparing the top 10 ranked features (Jaccard index). PanomiX consistently identified common features, indicating it effectively captures both linear and non-linear relationships in transcript–phenotype data while maintaining high predictive performance.

## Discussion

In this study we introduced panomiX, a user-friendly toolbox for multi-omics data integration. We demonstrated the versatility and robustness of panomiX on a unique multi-omic data set designed to identify transcriptomic and metabolic features associated with phenotypic traits. By addressing key challenges in normalization, variance analysis, and predictive modeling, we showed how panomiX enables extracting biologically meaningful insights from such complex datasets.

Our tool provided a robust workflow for transcriptomics and FTIR data, including tailored normalization and scaling of individual datasets. As shown in **Fig. 3**, stepwise transformations - here, from raw FTIR spectra to normalized outputs - improve data quality and interpretability through baseline correction and smoothing. This mitigates biases in raw datasets and ensures compatibility with downstream analyses such as PCA. PanomiX’s PCA and ANOVA functions further standardize and visualize variance across separate and combined datasets, revealing consistent clustering patterns driven by time points and treatment conditions. These insights highlight the complementary nature of multiple omics layers, setting the stage for more targeted multi-omics analyses.

PanomiX implementation of XGBoost consistently outperformed sPLS for the integration of our tomato heat stress datasets, showing higher accuracy, robustness, and feature stability. Its XGBoost-based framework effectively handles varying collinearity and identifies non-linear relationships, a key limitation in linear models. High Jaccard indices confirm feature stability across correlation gradients, while GO enrichment validates the biological relevance of identified predictors. Notably, the differential performance between control and treatment models underscores condition-specific mechanisms, exemplified by transcripts such as cytochrome b559 subunit alpha and phylloplanin in our experiment. These results highlight panomiX’s capacity to reveal critical molecular interactions and support hypothesis-driven research.

By integrating transcriptomics, FTIR, and phenomics data, panomiX uncovers shared predictors - such as cytochrome b559 subunit alpha and specific FTIR spectra - indicating convergent biological mechanisms. These cross-layer insights reinforce panomiX’s ability to reveal broadly relevant molecular features. Moreover, the option to add user-defined elements (e.g., heat-stress genes) illustrates its versatility for hypothesis-driven analyses. Identified interactions among kinases and stress-response transcripts emphasize the tool’s value for mapping complex biological networks, and point to the need for evaluating both positive and negative feature associations when predicting phenotypic outcomes.

While panomiX offers significant advancements, several areas for improvement are worth consideration. Currently, normalization capabilities are limited to transcriptomics and FTIR data, leaving scope for extending these features to other omics layers, such as proteomics and metabolomics. Expanding the curated feature databases and incorporating pathway-level analyses could provide deeper insights into the functional implications of predictive features.

## Code and Data Availability

The code for panomiX is freely available at https://github.com/NAMlab/panomiX-tool under the terms of the MIT license (also archived at Zenodo at time of publication: https://doi.org/10.5281/zenodo.15193421). The sequence data for this study have been deposited in the European Nucleotide Archive (ENA) at EMBL-EBI under accession number PRJEB85881 (https://www.ebi.ac.uk/ena/browser/view/PRJEB85881). Phenotyping and FTIR data as well as pre-processed inputs for reproducing the results of this article with panomiX are available at https://doi.org/10.5447/ipk/2025/3.

## Acknowledgements

We gratefully acknowledge the expert technical assistance of Annett Berge and Andrea Apelt during sample processing and molecular assay preparation and of Jacqueline Pohl during Illumina library preparation and RNA sequencing, as well as Gunda Wehrstedt and Ingo Mücke for plant cultivation and phenotyping and Steffen Wagner for experimental support in the FTIR analysis. We also thank Anne Fiebig and Clemens Porsche for their support in submitting the sequencing data to ENA and Daniel Arend and Matthias Lange for publishing the phenotyping and FTIR data on e!DAL.

## Author contributions

A.S., J.S., D.P., K.P., and N.T. wrote the manuscript with input from all the authors. D.P. planned, conducted, and coordinated the experimental work for transcriptomics analysis, FTIR analysis, and phenotyping under J.S. and K.N. supervision. A.H. performed RNA sequencing, H.R., and L.B. performed FTIR analysis. A.S. developed the toolbox under J.S.’s supervision. A.S., and D.P. performed the data analysis and machine learning modeling under J.S.’s supervision.

## Supplementary Materials

Table S1. The correlated matrix of all the 138 phenotypes we obtained from the IPK Phenosphere platform reduced to 27 uncorrelated phenotype clusters.

Table S2. Transcripts and FTIR features were consistently identified as top predictors across all three omics-specific models.

Table S3. Transcripts showed positive interactions with both the predicted features and the target phenotypes (scenario a).

Table S4. Transcripts showed interactions with both the predicted features and the target phenotypes from no constraints, random constraints, and monotonics constraints model (scenario b).

**Fig. S1.**
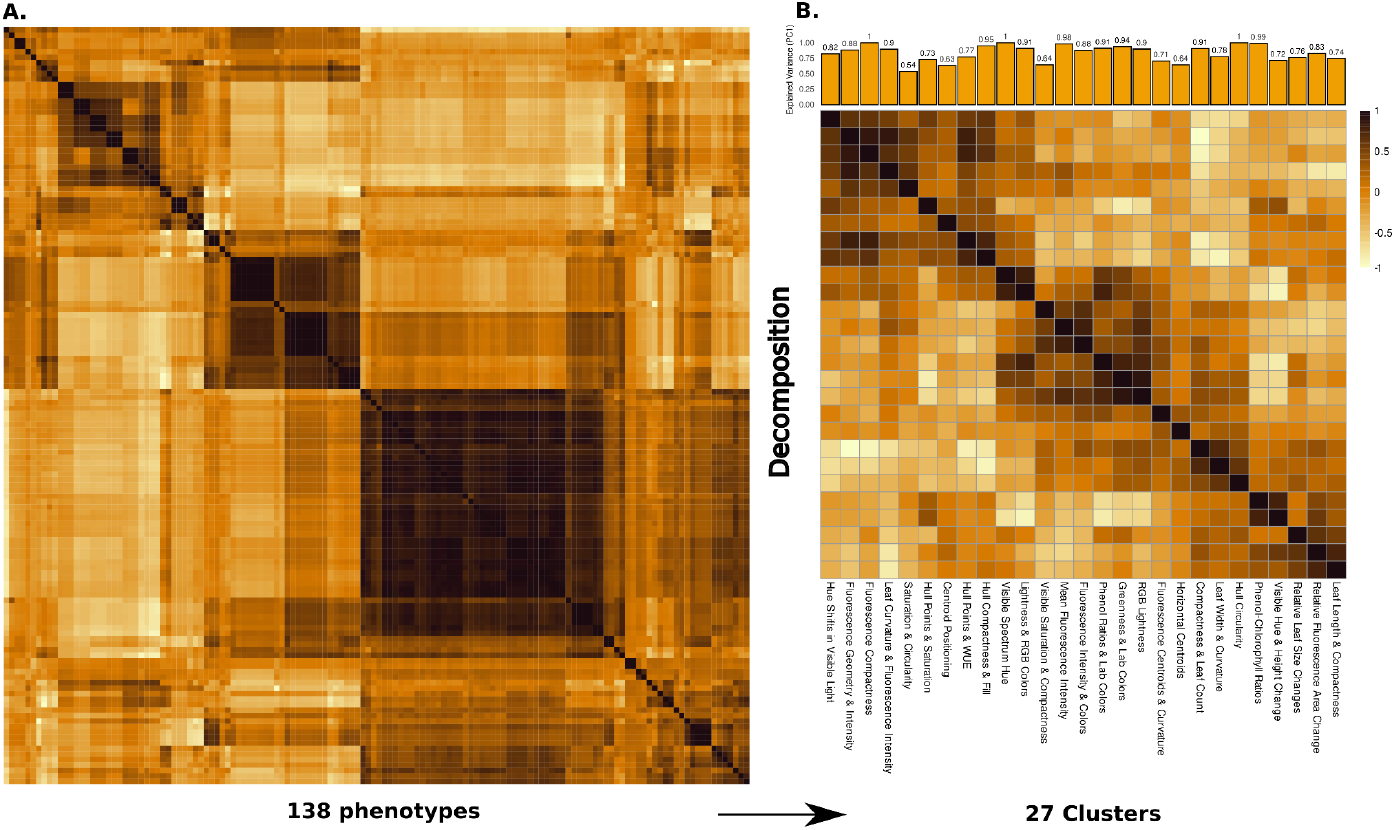
(A) Correlated matrix of all the 138 phenotypes we obtained from the IPK Phenosphere platform. (B) Here we have obtained the cluster of correlated phenotypes using hierarchical clustering and then we used PCA based decomposition on each cluster. Total explained variance (for PC1) for each dataset has been illustrated in the top of the figure (B). On the bottom we represented the correlation plot of each decomposed phenotype cluster. More info (Table S1)

**Fig. S2.**
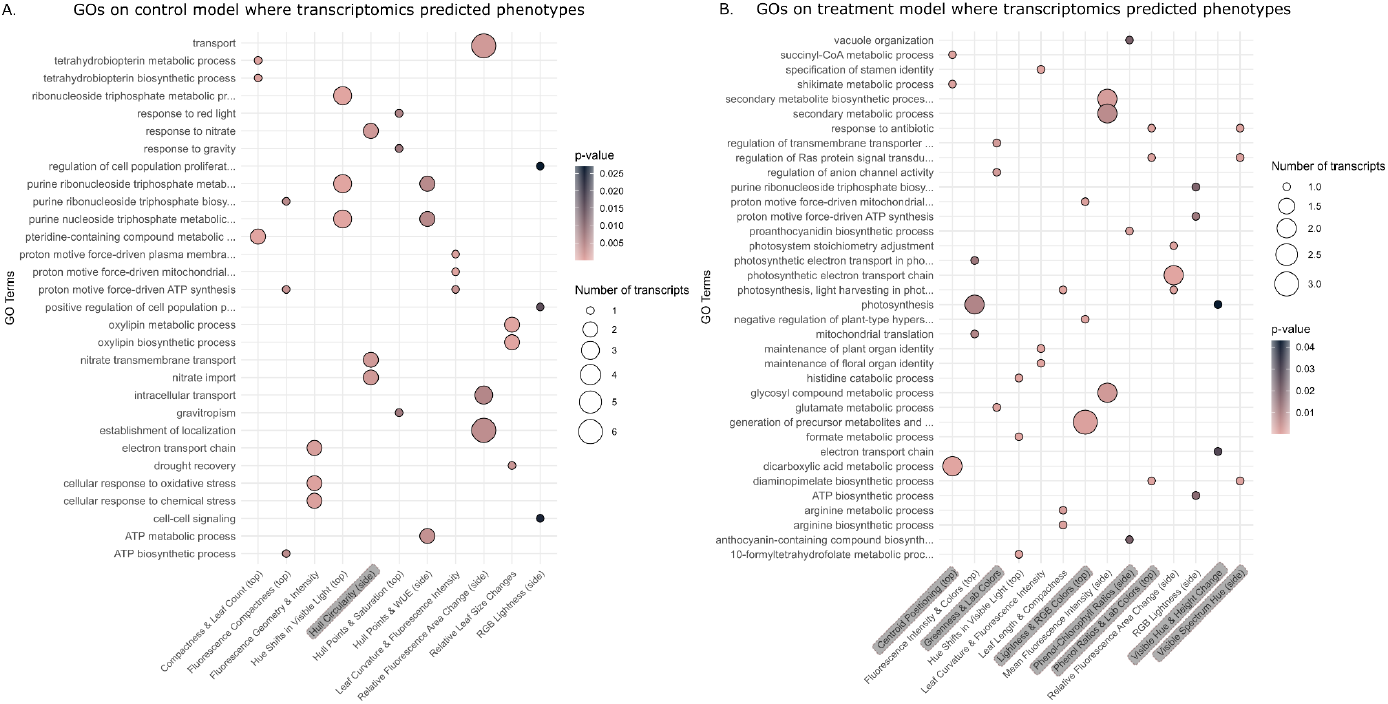
(A) Top 3 gene ontology (biological processes) for the features selected from the control model (transcriptomics-predicting phenotypes) (B) Top 3 gene ontology (biological processes) for the features selected from the treatment model (transcriptomics-predicting phenotypes). Models shown with grey shading have a predictive performance of R^2^ > 0.3 and replicate-model R^2^ > 0.5. Models without shading have R^2^ > 0.5 and replicate-model R^2^ > 0.7.

**Fig. S3.**
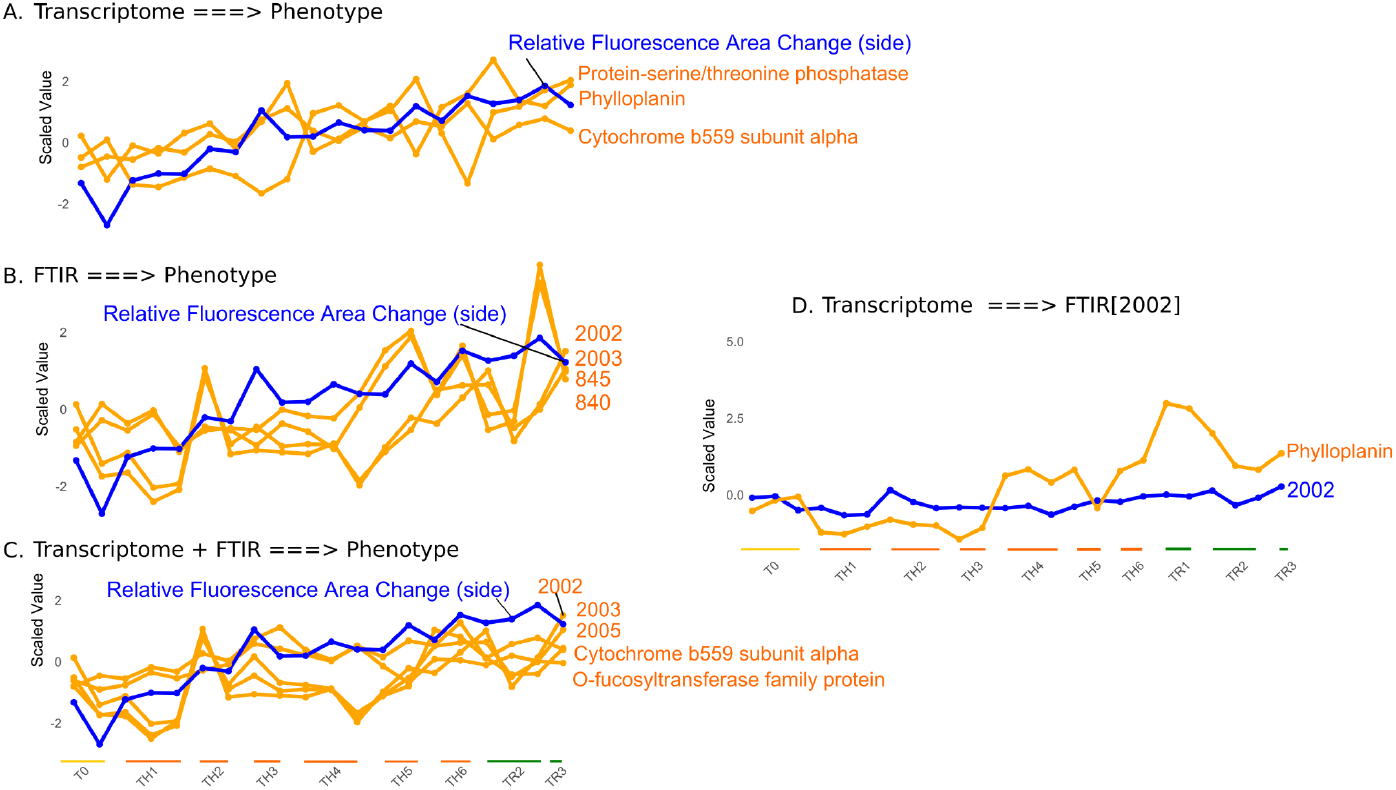
Features selected from different integrative models, where important features are highlighted in orange and the predicted variable is shown in blue. (A) Expression profile for the Transcriptome predicting Relative Fluorescence Area Change (side) (B) Feature important for the FTIR predicting Relative Fluorescence Area Change (side) model (C) Feature important for the Transcriptome + FTIR predicting Relative Fluorescence Area Change (side) model (D) Expression profile for the Transcriptome predicting FTIR wavelength [2002].

## References

[1] D. Houle, D. R. Govindaraju, and S. Omholt, “Phenomics: the next challenge,” Nat. Rev. Genet., vol. 11, no. 12, pp. 855–866, Dec. 2010, doi: 10.1038/nrg2897.

[2] R. Pieruschka and U. Schurr, “Plant Phenotyping: Past, Present, and Future,” Plant Phenomics, vol. 2019, p. 7507131, Mar. 2019, doi: 10.34133/2019/7507131.

[3] S. Ma et al., “WheatOmics: A platform combining multiple omics data to accelerate functional genomics studies in wheat,” Mol. Plant, vol. 14, no. 12, pp. 1965–1968, Dec. 2021, doi: 10.1016/j.molp.2021.10.006.

[4] W. Kong et al., “Genomic analysis of 1,325 Camellia accessions sheds light on agronomic and metabolic traits for tea plant improvement,” Nat. Genet., pp. 1–11, Mar. 2025, doi: 10.1038/s41588-025-02135-z.

[5] J. Szymański et al., “Analysis of wild tomato introgression lines elucidates the genetic basis of transcriptome and metabolome variation underlying fruit traits and pathogen response,” Nat. Genet., vol. 52, no. 10, pp. 1111–1121, Oct. 2020, doi: 10.1038/s41588-020-0690-6.

[6] J. Chen, Q. Li, and D. Jiang, “From Images to Loci: Applying 3D Deep Learning to Enable Multivariate and Multitemporal Digital Phenotyping and Mapping the Genetics Underlying Nitrogen Use Efficiency in Wheat,” Plant Phenomics, vol. 6, p. 0270, Dec. 2024, doi: 10.34133/plantphenomics.0270.

[7] Q. Lou et al., “Phenomics-assisted genetic dissection and molecular design of drought resistance in rice,” Plant Commun., vol. 6, no. 3, Mar. 2025, doi: 10.1016/j.xplc.2024.101218.

[8] C. Fu et al., “Transcriptomic and methylomic analyses provide insights into the molecular mechanism and prediction of heterosis in rice,” Plant J., vol. 115, no. 1, pp. 139–154, 2023, doi: 10.1111/tpj.16217.

[9] T. B. Mersha, “From Mendel to multi-omics: shifting paradigms,” Eur. J. Hum. Genet., vol. 32, no. 2, pp. 139–142, Feb. 2024, doi: 10.1038/s41431-023-01420-x.

[10] D. Knoch et al., “Integrated multi-omics analyses and genome-wide association studies reveal prime candidate genes of metabolic and vegetative growth variation in canola,” Plant J., vol. 117, no. 3, pp. 713–728, 2024, doi: 10.1111/tpj.16524.

[11] D. Feldner-Busztin et al., “Dealing with dimensionality: the application of machine learning to multi-omics data,” Bioinformatics, vol. 39, no. 2, p. btad021, 2023.

[12] Y. Luo, C. Zhao, and F. Chen, “Multiomics Research: Principles and Challenges in Integrated Analysis,” BioDesign Res., vol. 6, p. 0059, Jan. 2024, doi: 10.34133/bdr.0059.

[13] B. Wang et al., “Similarity network fusion for aggregating data types on a genomic scale,” Nat. Methods, vol. 11, no. 3, pp. 333–337, 2014.

[14] C. Dimitrakopoulos et al., “Network-based integration of multi-omics data for prioritizing cancer genes,” Bioinformatics, vol. 34, no. 14, pp. 2441–2448, 2018.

[15] C. J. Vaske et al., “Inference of patient-specific pathway activities from multi-dimensional cancer genomics data using PARADIGM,” Bioinformatics, vol. 26, no. 12, pp. i237–i245, 2010.

[16] R. Louhimo and S. Hautaniemi, “CNAmet: an R package for integrating copy number, methylation and expression data,” Bioinformatics, vol. 27, no. 6, pp. 887–888, 2011.

[17] H. Nguyen, S. Shrestha, S. Draghici, and T. Nguyen, “PINSPlus: a tool for tumor subtype discovery in integrated genomic data,” Bioinformatics, vol. 35, no. 16, pp. 2843–2846, 2019.

[18] N. Rappoport and R. Shamir, “NEMO: cancer subtyping by integration of partial multi-omic data,” Bioinformatics, vol. 35, no. 18, pp. 3348–3356, 2019.

[19] A. Orlichenko et al., “Latent Similarity Identifies Important Functional Connections for Phenotype Prediction,” IEEE Trans. Biomed. Eng., vol. 70, no. 6, pp. 1979–1989, Jun. 2023, doi: 10.1109/TBME.2022.3232964.

[20] R. Argelaguet et al., “Multi‐Omics Factor Analysis—a framework for unsupervised integration of multi‐omics data sets,” Mol. Syst. Biol., vol. 14, no. 6, p. e8124, Jun. 2018, doi: 10.15252/msb.20178124.

[21] R. Shen et al., “Integrative subtype discovery in glioblastoma using iCluster,” PloS One, vol. 7, no. 4, p. e35236, 2012.

[22] P. Ray, L. Zheng, J. Lucas, and L. Carin, “Bayesian joint analysis of heterogeneous genomics data,” Bioinformatics, vol. 30, no. 10, pp. 1370–1376, 2014.

[23] C. Zhang, Y. Chen, T. Zeng, C. Zhang, and L. Chen, “Deep latent space fusion for adaptive representation of heterogeneous multi-omics data,” Brief. Bioinform., vol. 23, no. 2, p. bbab600, Mar. 2022, doi: 10.1093/bib/bbab600.

[24] C. Meng, D. Helm, M. Frejno, and B. Kuster, “moCluster: Identifying Joint Patterns Across Multiple Omics Data Sets,” J. Proteome Res., vol. 15, no. 3, pp. 755–765, Mar. 2016, doi: 10.1021/acs.jproteome.5b00824.

[25] F. Rohart, B. Gautier, A. Singh, and K.-A. Lê Cao, “mixOmics: An R package for ‘omics feature selection and multiple data integration,” PLoS Comput. Biol., vol. 13, no. 11, p. e1005752, 2017.

[26] K. Kaitoh and Y. Yamanishi, “TRIOMPHE: Transcriptome-Based Inference and Generation of Molecules with Desired Phenotypes by Machine Learning,” J. Chem. Inf. Model., vol. 61, no. 9, pp. 4303–4320, Sep. 2021, doi: 10.1021/acs.jcim.1c00967.

[27] T. Chen and C. Guestrin, “XGBoost: A Scalable Tree Boosting System,” in Proceedings of the 22nd ACM SIGKDD International Conference on Knowledge Discovery and Data Mining, San Francisco California USA: ACM, Aug. 2016, pp. 785–794. doi: 10.1145/2939672.2939785.

[28] H. Chen, I. C. Covert, S. M. Lundberg, and S.-I. Lee, “Algorithms to estimate Shapley value feature attributions,” Nat. Mach. Intell., vol. 5, no. 6, pp. 590–601, 2023.

[29] D. Camejo, P. Rodríguez, M. Angeles Morales, J. Miguel Dell’Amico, A. Torrecillas, and J. J. Alarcón, “High temperature effects on photosynthetic activity of two tomato cultivars with different heat susceptibility,” J. Plant Physiol., vol. 162, no. 3, pp. 281–289, Mar. 2005, doi: 10.1016/j.jplph.2004.07.014.

[30] C. Bita and T. Gerats, “Plant tolerance to high temperature in a changing environment: scientific fundamentals and production of heat stress-tolerant crops,” Front. Plant Sci., vol. 4, Jul. 2013, doi: 10.3389/fpls.2013.00273.

[31] M. M. Raja et al., “Pollen development and function under heat stress: from effects to responses,” Acta Physiol. Plant., vol. 41, no. 4, p. 47, Mar. 2019, doi: 10.1007/s11738-019-2835-8.

[32] C. Jonak, S. Kiegerl, W. Ligterink, P. J. Barker, N. S. Huskisson, and H. Hirt, “Stress signaling in plants: a mitogen-activated protein kinase pathway is activated by cold and drought.,” Proc. Natl. Acad. Sci., vol. 93, no. 20, pp. 11274–11279, Oct. 1996, doi: 10.1073/pnas.93.20.11274.

[33] Y.-X. Lin, H.-Y. Jiang, Z.-X. Chu, X.-L. Tang, S.-W. Zhu, and B.-J. Cheng, “Genome-wide identification, classification and analysis of heat shock transcription factor family in maize,” BMC Genomics, vol. 12, no. 1, p. 76, Jan. 2011, doi: 10.1186/1471-2164-12-76.

[34] H. Tanaka et al., “Abiotic stress-inducible receptor-like kinases negatively control ABA signaling in Arabidopsis,” Plant J., vol. 70, no. 4, pp. 599–613, 2012, doi: 10.1111/j.1365-313X.2012.04901.x.

[35] Y. Liu, Z. Feng, W. Zhu, J. Liu, and Y. Zhang, “Genome-Wide Identification and Characterization of Cysteine-Rich Receptor-Like Protein Kinase Genes in Tomato and Their Expression Profile in Response to Heat Stress,” Diversity, vol. 13, no. 6, Art. no. 6, Jun. 2021, doi: 10.3390/d13060258.

[36] S. Mo et al., “Mitogen-activated protein kinase action in plant response to high-temperature stress: a mini review,” Protoplasma, vol. 258, no. 3, pp. 477–482, May 2021, doi: 10.1007/s00709-020-01603-z.

[37] M. Pardo-Hernández, P. García-Pérez, L. Lucini, and R. M. Rivero, “Multi-omics exploration of the involvement of ABA in identifying unique molecular markers for single and combined stresses in tomato plants,” J. Exp. Bot., p. erae372, Sep. 2024, doi: 10.1093/jxb/erae372.

[38] M. I. Love, W. Huber, and S. Anders, “Moderated estimation of fold change and dispersion for RNA-seq data with DESeq2,” Genome Biol., vol. 15, no. 12, p. 550, Dec. 2014, doi: 10.1186/s13059-014-0550-8.

[39] K. H. Liland and B.-H. Mevik, “baseline: Baseline Correction of Spectra.” p. 1.3-5, Jan. 10, 2011. doi: 10.32614/CRAN.package.baseline.

[40] U. Ligges, T. Short, and P. Kienzle, “signal: Signal Processing.” p. 1.8-1, Dec. 10, 2006. doi: 10.32614/CRAN.package.signal.

[41] M. U. A. Bromba and Horst. Ziegler, “Application hints for Savitzky-Golay digital smoothing filters,” Anal. Chem., vol. 53, no. 11, pp. 1583–1586, Sep. 1981, doi: 10.1021/ac00234a011.

[42] J. Baglama, L. Reichel, and B. W. Lewis, “irlba: Fast Truncated Singular Value Decomposition and Principal Components Analysis for Large Dense and Sparse Matrices.” p. 2.3.5.1, May 27, 2011. doi: 10.32614/CRAN.package.irlba.

[43] M. Kuhn, “caret: Classification and Regression Training.” p. 7.0–1, Oct. 05, 2007. doi: 10.32614/CRAN.package.caret.

[44] M. B. Kursa and W. R. Rudnicki, “Feature selection with the Boruta package,” J. Stat. Softw., vol. 36, pp. 1–13, 2010.

[45] D. Attali, “shinyjs: Easily Improve the User Experience of Your Shiny Apps in Seconds.” p. 2.1.0, Apr. 22, 2015. doi: 10.32614/CRAN.package.shinyjs.

[46] C. Sievert et al., “plotly: Create Interactive Web Graphics via ‘plotly.js.’” p. 4.10.4, Nov. 17, 2015. doi: 10.32614/CRAN.package.plotly.

[47] Y. Xie, J. Cheng, and X. Tan, “DT: A Wrapper of the JavaScript Library ‘DataTables.’” p. 0.33, Jun. 09, 2015. doi: 10.32614/CRAN.package.DT.

[48] D. Attali and A. Sali, “shinycssloaders: Add Loading Animations to a ‘shiny’ Output While It’s Recalculating.” p. 1.1.0, May 10, 2017. doi: 10.32614/CRAN.package.shinycssloaders.

[49] C. Sievert, J. Cheng, and G. Aden-Buie, “bslib: Custom ‘Bootstrap’ ‘Sass’ Themes for ‘shiny’ and ‘rmarkdown.’” p. 0.9.0, Jan. 25, 2021. doi: 10.32614/CRAN.package.bslib.

[50] C. Sievert, “bsicons: Easily Work with ‘Bootstrap’ Icons.” p. 0.1.2, Nov. 22, 2022. doi: 10.32614/CRAN.package.bsicons.

[51] R. Ihaka and R. and Gentleman, “R: A Language for Data Analysis and Graphics,” J. Comput. Graph. Stat., vol. 5, no. 3, pp. 299–314, Sep. 1996, doi: 10.1080/10618600.1996.10474713.

[52] W. Chang et al., “shiny: Web Application Framework for R.” p. 1.10.0, Dec. 01, 2012. doi: 10.32614/CRAN.package.shiny.

[53] A. Atkins, T. Allen, H. Wickham, J. McPherson, and J. Allaire, “rsconnect: Deploy Docs, Apps, and APIs to ‘Posit Connect’, ‘shinyapps.io’, and ‘RPubs.’” p. 1.3.4, Mar. 20, 2016. doi: 10.32614/CRAN.package.rsconnect.

[54] C. Klukas, D. Chen, and J.-M. Pape, “Integrated Analysis Platform: An Open-Source Information System for High-Throughput Plant Phenotyping,” Plant Physiol., vol. 165, no. 2, pp. 506–518, Jun. 2014, doi: 10.1104/pp.113.233932.

[55] D. Arend et al., “Quantitative monitoring of Arabidopsis thaliana growth and development using high-throughput plant phenotyping,” Sci. Data, vol. 3, no. 1, p. 160055, Aug. 2016, doi: 10.1038/sdata.2016.55.

[56] N. L. Bray, H. Pimentel, P. Melsted, and L. Pachter, “Near-optimal probabilistic RNA-seq quantification,” Nat. Biotechnol., vol. 34, no. 5, pp. 525–527, May 2016, doi: 10.1038/nbt.3519.

[57] D. Psaroudakis et al., “Identification of core, conditional and crosstalk components of tomato heat stress response using integrative transcriptomics and orthology,” 2024, Accessed: Jan. 08, 2025. [Online]. Available: https://www.researchsquare.com/article/rs-4337825/latest

[58] A. Guendel, A. Hilo, H. Rolletschek, and L. Borisjuk, “Probing the Metabolic Landscape of Plant Vascular Bundles by Infrared Fingerprint Analysis, Imaging and Mass Spectrometry,” Biomolecules, vol. 11, no. 11, Art. no. 11, Nov. 2021, doi: 10.3390/biom11111717.

[59] S. Zarbakhsh and A. R. Shahsavar, “Exogenous γ-aminobutyric acid improves the photosynthesis efficiency, soluble sugar contents, and mineral nutrients in pomegranate plants exposed to drought, salinity, and drought-salinity stresses,” BMC Plant Biol., vol. 23, no. 1, p. 543, Nov. 2023, doi: 10.1186/s12870-023-04568-2.

[60] S. Singh et al., “Identification of key genes and molecular pathways regulating heat stress tolerance in pearl millet to sustain productivity in challenging ecologies,” Front. Plant Sci., vol. 15, p. 1443681, 2024.

[61] M. Koul, L. Thomas, and K. Karmakar, “Functional aspects of solanaceae trichomes in heavy metal detoxification,” Nord. J. Bot., vol. 39, no. 5, p. njb.03171, May 2021, doi: 10.1111/njb.03171.

[62] Y. Zheng et al., “iTAK: a program for genome-wide prediction and classification of plant transcription factors, transcriptional regulators, and protein kinases,” Mol. Plant, vol. 9, no. 12, pp. 1667–1670, 2016.

[63] H. Rahman, X.-Y. Wang, Y.-P. Xu, Y.-H. He, and X.-Z. Cai, “Characterization of tomato protein kinases embedding guanylate cyclase catalytic center motif,” Sci. Rep., vol. 10, no. 1, p. 4078, 2020.

[64] S. Graci and A. Barone, “Tomato plant response to heat stress: a focus on candidate genes for yield-related traits,” Front. Plant Sci., vol. 14, p. 1245661, 2024.

[65] V. Berková et al., “The fungus Acremonium alternatum enhances salt stress tolerance by regulating host redox homeostasis and phytohormone signaling,” Physiol. Plant., vol. 176, no. 3, p. e14328, May 2024, doi: 10.1111/ppl.14328.

[66] Y. Zhao et al., “The calcium‐dependent protein kinase ZmCDPK7 functions in heat‐stress tolerance in maize,” J. Integr. Plant Biol., vol. 63, no. 3, pp. 510–527, Mar. 2021, doi: 10.1111/jipb.13056.

[67] J. Zhang et al., “Releasing a sugar brake generates sweeter tomato without yield penalty,” Nature, pp. 1–10, 2024.

